# HIV Diversity Considerations in the Application of the Intact Proviral DNA Assay (IPDA)

**DOI:** 10.1101/2020.05.26.115006

**Authors:** Natalie N. Kinloch, Yanqin Ren, Winiffer D. Conce Alberto, Winnie Dong, Pragya Khadka, Szu Han Huang, Talia M. Mota, Andrew Wilson, Aniqa Shahid, Don Kirkby, Marianne Harris, Colin Kovacs, Erika Benko, Mario A. Ostrowski, Perla M. Del Rio Estrada, Avery Wimpelberg, Christopher Cannon, W. David Hardy, Lynsay MacLaren, Harris Goldstein, Chanson J. Brumme, Guinevere Q. Lee, Rebecca M. Lynch, Zabrina L. Brumme, R. Brad Jones

**Author notes:** These authors contributed equally (Natalie N. Kinloch, Yanqin Ren). These are co-senior/co-corresponding authors.

## Abstract

The Intact Proviral DNA Assay (IPDA) was developed to address the critical need for a precise and scalable method for intact HIV reservoir quantification^1^. This duplexed droplet digital PCR (ddPCR) assay simultaneously targets the HIV Packaging Signal (Ψ) and the Rev Responsive Element (RRE) within Envelope (*env*) to distinguish genomically intact proviruses against a large background of defective ones^2^. The IPDA requires less time, resources, and biological material than the gold standard for replication-competent HIV reservoir measurement, the Quantitative Viral Outgrowth Assay (QVOA)^3^, and is being adopted in research and clinical studies^4–7^. In our cohort of HIV-1 subtype B-infected individuals from North America however, the IPDA yielded a 28% failure rate due to HIV polymorphism. We further demonstrate that within-host HIV diversity can lead the IPDA to underestimate intact HIV reservoir size, which could negatively impact clinical trial results interpretation. While the IPDA represents an important methodological advance, HIV diversity should be addressed before its widespread adoption.

---

We applied the IPDA to 46 HIV subtype B-infected, virally-suppressed individuals from North America, yielding a median of 29 (interquartile range [IQR] 0-93) intact proviruses/million CD4+ T-cells (Extended Data Figure 1). Of note, the IPDA did not detect any intact (*i.e*. Ψ and *env* double-positive) proviruses in 17 participants (37%). In four of these individuals, both Ψ- and *env*-single-positive proviruses were detected, suggesting a true negative result (Extended Data Figure 3) In the remaining 13 individuals however, the IPDA did not detect Ψ- and/or *env*-single-positive proviruses above background levels despite recovery of replication competent HIV in 11/11 cases where QVOA was performed, suggesting that the assay failed to detect autologous proviruses. Specifically, in eight of these participants only Ψ-positive proviruses were detected, in four only *env*-positive proviruses were detected, and in one participant no proviruses were detected (Figure 1B; Extended Data Figure 3). This non-detection rate is consistent with Gaebler *et al* who reported a lack of detectable Ψ and *env* double-positive proviruses in 2/6 individuals (33%) using a quadruplexed qPCR assay that employs the IPDA probes^8^. Near-full-length single-genome proviral sequencing performed on IPDA *env*-negative participant BC-004 revealed mismatches to the *env* probe, where *in silico-*predicted reservoir distributions that took these polymorphisms into account (Figure 1C, left) differed markedly from the experimentally-obtained result (center). Substituting an autologous *env* probe rescued detection to *in silico*-predicted levels (Figure 1C, right), confirming that HIV polymorphism can cause the IPDA to fail. HIV sequencing revealed mismatches in the probe and/or at the 3’ end of a primer in all cases of presumed assay detection failure (Extended Data Figure 4), yielding an overall estimated failure rate of 13/46 (28%). Indeed, exclusion of these 13 datapoints markedly improved the correlation between IPDA and QVOA results among the subset of participants for whom sufficient biological material was available to perform the latter, from ρ= 0.03, p=0.83 to ρ= 0.35, p=0.08 (Figure 1C), bringing this correlation more in line with that reported by the original authors^1^. The correlation between total HIV *gag* DNA and QVOA by contrast was significant without exclusion of these datapoints (Extended Data Figure 2). Our findings thus reinforce the value of the IPDA but underscore the need for strategies to overcome HIV diversity.

**Figure 1:**
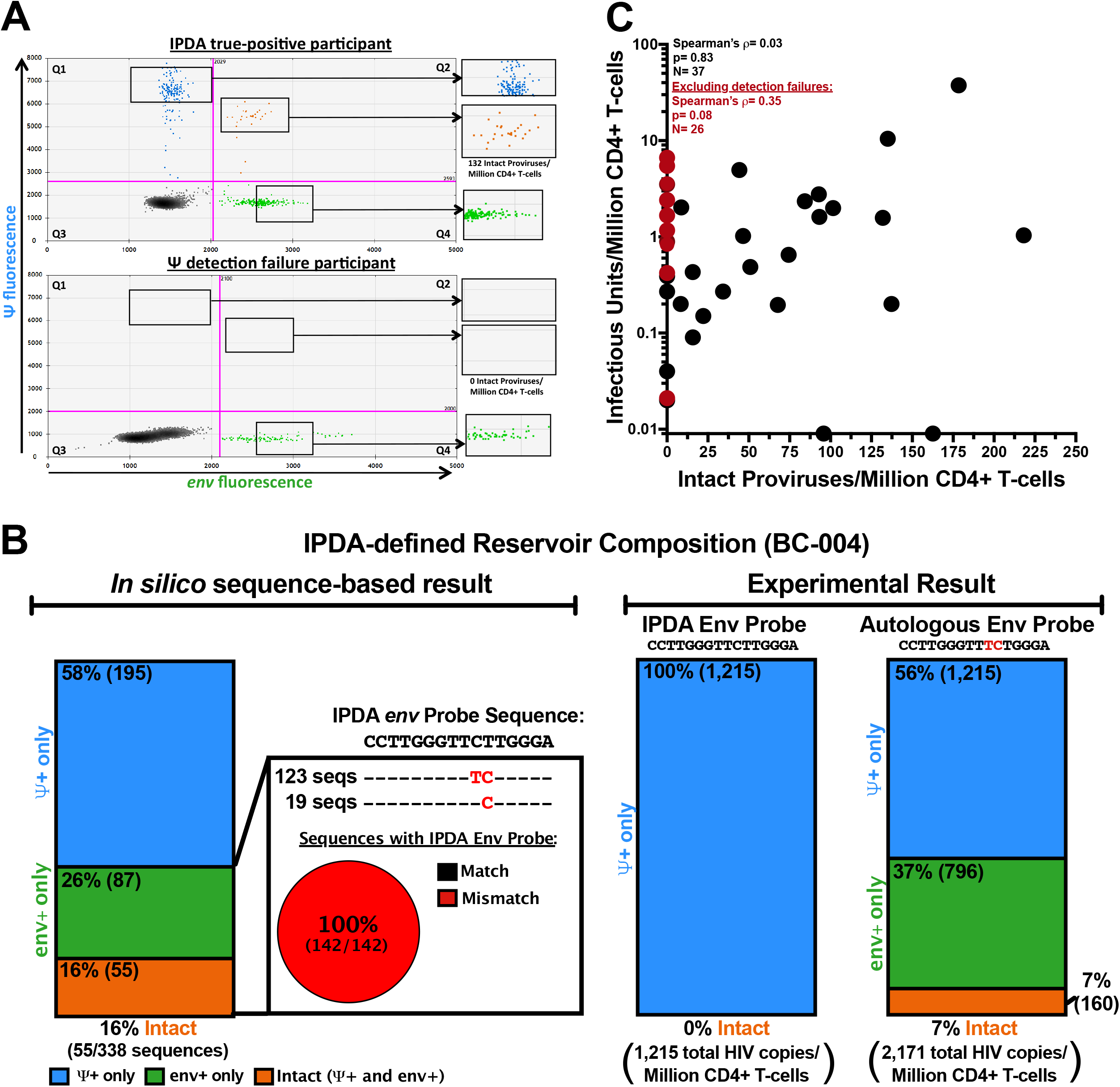
Inter-individual HIV diversity can lead to detection failure by the IPDA. **(A) Example IPDA 2D plots contrasting a true positive result from a case of detection failure.** 2D ddPCR plots showing Ψ-single positive events (Q1, blue), Ψ- and *env*-double positive events (Q2, orange), double-negative events (Q3, grey) and *env*-single positive events (Q4, green), for an IPDA true-positive individual (top) and a presumed case of Ψ detection failure (bottom). **(B) Inter-individual HIV diversity can lead to detection failure in the IPDA.** (**left**) Analysis of 338 near-full length proviruses from participant BC-004, who originally failed detection in *env*, revealed single nucleotide mismatches (red, inset) to the IPDA *env* probe. These sequences yielded *in silico* Ψ+ (blue), *env*+ (green) and intact (Ψ+*env*+; orange) provirus frequencies as shown. (**right**) Experimental results obtained with the published IPDA and autologous *env* probe, respectively. **(C) IPDA detection failures obscure correlation between QVOA and IPDA in a North American cohort.** Spearman’s correlation between reservoir size as measured by IPDA (Intact Proviruses/Million CD4+ T-cells) and QVOA (Infectious Units per Million CD4+ T-cells) in 37 virally-suppressed participants for whom the latter measurement was available (ρ= 0.03, p=0.83) and excluding presumed instances of IPDA detection failure (red datapoints, n= 26 remaining, ρ= 0.35, p=0.08). Two individuals for whom no replication competent viruses were detected (IUPM= 0) are plotted on the X-axis.

Although the original study reported no detection failures due to HIV polymorphism, the authors acknowledged that triage primer/probe sets would be required in such cases^1^. A follow-up study similarly acknowledged HIV diversity as a potential limitation, but reported no participants (of n = 81) for whom detection consistently failed for either Ψ or *env* amplicons^5^. By contrast, our results indicate that, as the IPDA is applied to diverse cohorts^4,5,7^, detection failures will occur, and not infrequently. The first step towards mitigation is awareness: samples that yield no Ψ or *env* single-positive proviruses should be flagged as ‘unreportable’ until HIV polymorphism has been addressed (*e.g*. using autologous primers/probes^1,5^). Analysis of HIV subtype B sequences from unique individuals in the Los Alamos HIV database revealed that 23% of 9,360 *env* sequences harbored at least one *env* probe mismatch (which is similar to our *env* detection failure rate of 9/46 or 20%), while 50% of 1,489 sequences harbored at least one Ψ probe mismatch (which is substantially greater than our Ψ detection failure rate of 5/46 or 11%). This suggests that the *env* reaction may be particularly sensitive to polymorphism.

Though laborious to correct, complete detection failures due to HIV polymorphism are nevertheless easy to flag. In contrast, within-host HIV diversity (see Figure 1C for an example) could lead the IPDA to underestimate intact reservoir size if the within-host variants were differentially detectable by the assay. Such partial detection failures would *not* be easy to identify. Moreover, if IPDA-detectable and non-detectable reservoir subpopulations were differentially susceptible to HIV cure interventions, this could lead to erroneous conclusions regarding intervention efficacy.

We illustrate this using HIV-specific broadly neutralizing antibodies (bNAbs). bNAbs can facilitate elimination of HIV-infected cells^9^ in part by targeting them for antibody-dependent cellular cytotoxicity (ADCC)^9,10^, and are being evaluated in clinical trials, some of which use the IPDA as a readout (*e.g*: ACTG A5386^6^). Participant 91C33 from a published trial^11^ provides a hypothetical example. This individual did not respond to (off-ART) infusions of the bNAbs 3BNC117 and 10-1074 because they harbored a plasma HIV subpopulation that was resistant to both bNAbs^11^ (Figure 2A). Of note, this plasma HIV subpopulation also harbored a mismatch to the IPDA *env* probe. We confirmed that the published IPDA could not detect templates harboring this mismatch, while those representing the bNAb-sensitive strains were detected readily (Figure 2A; Extended Data Figure 5). If a person harboring such diversity in their reservoir were to be successfully treated with one or both bNAbs, the IPDA could over-estimate the intervention’s effect (Figure 2C).

**Figure 2:**
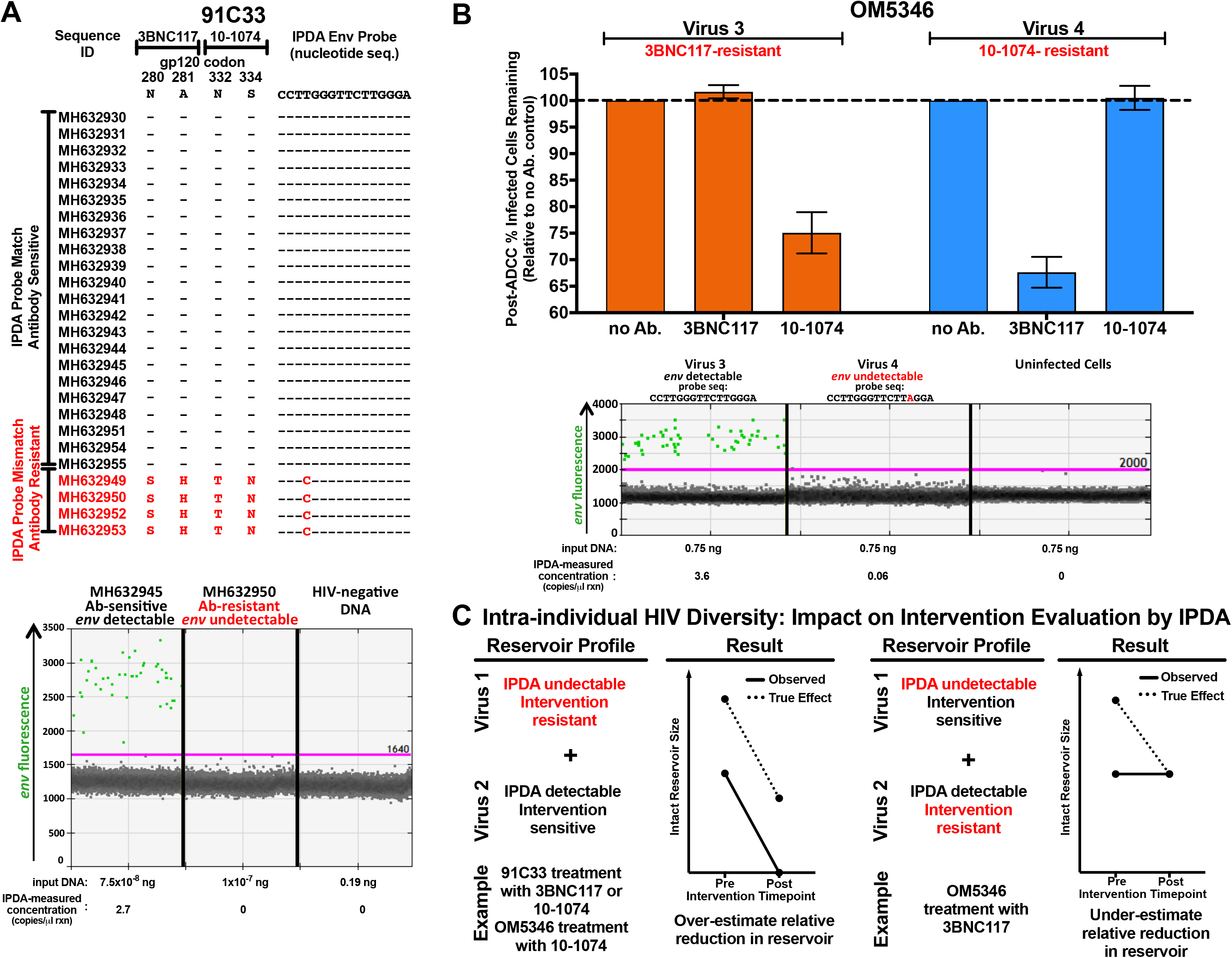
Intra-individual HIV diversity can lead to reservoir underestimation by IPDA. **(A) Example of heterogeneous within-host HIV variants that are differentially detectable by IPDA and differentially sensitive to bNAbs. (top)** Participant 91C33 harbored two major plasma HIV subpopulations: one that matched the IPDA *env* probe and was sensitive to bNAbs 3BNC117 and 10-1074, and the other that harbored a mismatch at position 4 of the IPDA *env* probe (red) and was resistant to 3BNC117 and 10-1074. Sequences are identified by their GenBank accession numbers; hyphens **(–)** indicate matches to the nucleotide or amino acid reference; red letters denote mismatches (IPDA *env* probe) or amino acid substitutions (bNAb). **(bottom)** Representative ddPCR *env* plots for MH632945 and MH632950; positive droplets are green and negative droplets are grey. Templates were purified *env* PCR products of equal length and comparable quantities. HIV-negative donor DNA served as a negative control. See Extended data Figure 5 for additional experiments. **(B) Example of heterogeneous within-host HIV variants that are differentially detectable by IPDA and differentially sensitive to bNAb-mediated ADCC. (top)** Two viruses (“virus 3” and “virus 4”) isolated from participant OM5346’s reservoir exhibited different sensitivities to bNAb-mediated ADCC: virus 3 was sensitive to 10-1074-mediated ADCC but resistant to 3BNC117, while virus 4 was sensitive to 3BNC117-mediated ADCC but resistant to 10-1074. Error bars indicate standard deviation from three technical replicates; see Extended Data Figure 7 for gating strategy and raw flow data. **(bottom)** Representative 1D IPDA *env* ddPCR plots for cells infected *in vitro* with virus 3, which matched the IPDA *env* probe sequence, and virus 4, which harbored the G13A mismatch. Uninfected cells served as a negative control. Both viruses were readily detectable using an alternative primer/probe set (see Extended data Figure 8A). **(C) Possible impacts of intra-individual HIV diversity on IPDA-measured changes in intact reservoir size following a hypothetical intervention.** (**left**) An intervention’s impact on the reservoir could be *overestimated* when a subset of within-host HIV sequences are undetectable by IPDA and *resistant* to the treatment (*e.g*. OM5346 virus 4 during treatment with 10-1074, or 91C33 during treatment with 3BNC117 or 10-1074). The solid line indicates the observed effect by IPDA; dashed line indicates the true effect. **(right)** An intervention’s impact on the reservoir could be *underestimated* when a subset of within-host HIV sequences are undetectable by IPDA and *sensitive* to treatment (*e.g*. OM5346 virus 4 during treatment with 3BNC117). In the worst-case scenario shown here, one could erroneously conclude that the treatment had no effect.

ARV-suppressed participant OM5346 provides another example. Pre-ART HIV drug resistance genotyping identified subtype B infection, however single-genome sequencing of pre-ART plasma and proviruses sampled during long-term ART revealed co-infection with a non-B strain (Extended Data Figure 6) that harbored a G-to-A mismatch at position 13 (G13A) of the *env* probe but no other critical mismatches to the IPDA primers. While the IPDA is only designed for subtype B, G13A is the most frequent *env* probe polymorphism [~5%] in subtype B (and was observed in 2/10 study participants for whom *env* was sequenced). Moreover, the original report indicated that the IPDA could detect G13A, at least when present on a plasmid template^1^. Using QVOA, we isolated replication-competent subtype B (“virus 3”) and coinfecting (“virus 4”) viruses from OM5346’s reservoir, confirming both of these as true intended IPDA targets (Figure 2B). We further observed that these viruses were differentially susceptible to 3BNC117- and 10-1074-mediated ADCC: while virus 3-infected cells could be eliminated by 10-1074-, but not 3BNC117-, mediated ADCC (25% and 0% relative reduction, respectively), the opposite was true for virus 4 (0% and 32% relative reduction, respectively) (Figure 2B, Extended Data Figure 7). At concentrations approximating participant samples, the IPDA could readily detect provirus 3, but could not distinguish provirus 4 (Figure 2B, Extended Data Figure 8A). Similarly, at participant-like concentrations, the IPDA could not detect a synthetic DNA template encoding the virus 4 sequence, though noticeable signal elevation occurred at ~100-fold higher template concentrations (Figure 2B, Extended Data Figure 8B). Importantly, a secondary *env* primer/probe set (see below and methods) readily detected both viruses at all concentrations, regardless of template type. If this individual were to be successfully treated with 10-1074 the IPDA would overestimate the intervention’s effect on the reservoir, whereas if the individual were to be successfully treated with 3BNC117, the IPDA would erroneously conclude that the intervention had no effect (Figure 2C). Our findings, taken together with Gaebler *et al’s* observation that 4 of 9 (44%) studied reservoirs were heterogeneous in an IPDA probe region^8^, suggests that the impact of within-host HIV diversity on IPDA accuracy may be non-negligible.

During our investigation we identified another source of potential error, resulting from variable spillover of fluorescence from the Ψ channel into the *env* channel, which has implications for the appropriate placement of thresholds defining negative and positive populations. Importantly, the extent of spillover is HIV sequence-specific. We demonstrate this by applying the IPDA to synthetic templates encoding the Ψ regions of OM5346 viruses 3 and 4, without a corresponding *env* template present (Extended Data Figure 9). Virus 3, which harbors mismatches to the Ψ probe, produced modest yet clearly discernible Ψ signal that did not spill over into the *env* channel. By contrast, virus 4 yielded high-amplitude Ψ signal that spilled over noticeably. Thus, drawing a tight threshold based on a template-negative (or virus 3-containing) sample and applying this threshold to virus 4 creates *env* signal when no *env* template was present. This Ψ channel spillover for virus 4 thus likely contributed to the small amount of ‘*env* signal’ in Fig. 2B (events adjoining the negative population). This observation underscores the instrument manufacturer’s instruction that appropriate thresholds can only be drawn if negative and positive populations clearly separate from one another.

The IPDA offers major scalability advantages over existing molecular^8^ or culture-based approaches^3^, and Gaebler *et al*. confirmed that, of any two probes, those of the IPDA offered the greatest selectivity for intact proviruses^8^. Any molecular assay targeting a genetically variable pathogen however must address polymorphism. While the requirement to discriminate proviral defects in the Ψ region limits the placement and sequence of this probe^2^, we developed a secondary primer/probe set in the intact-discriminating RRE region^1^, approximately 50 bases downstream of the original location, as a first step towards addressing diversity. This primer/probe set rescued detection of *env*-positive proviruses in 9/9 participants with IPDA *env* detection failure (Extended Data Figure 10A). When we applied it to 36 participants for whom the IPDA detected *env*-positive proviruses (excluding OM5346 who harbored within-host diversity), it failed to detect *env*-positive proviruses in 3 (8%) individuals, indicating that it is not a universal solution. For the remaining 33 (92%) however it yielded measurements that were not significantly different from the IPDA (Extended Data Figure 10B). The secondary primer/probe set could therefore be used to identify instances of detection failure, though users must note that it cannot discriminate hypermutated sequences.

We identify HIV diversity as a source of error in the IPDA. Within-host HIV diversity represents a challenge given the HIV reservoir’s dynamic nature^12^ and particularly when this diversity is linked to intervention susceptibility. Given the clear value of the IPDA, iterative efforts to refine this assay are a priority. In the meantime, proviral sequencing should be performed for clinical trial participants, and ‘backup’ and/or autologous primers/probes used to mitigate HIV diversity, an all-too-familiar challenge in viral quantification.

## Supplementary Information (Methods)

### Participants and Ethics Statement

We studied 46 individuals with HIV-1 subtype B infection who were recruited to cohorts in Toronto (N=19), Vancouver (N=15), New York City (N=4), Washington, D.C. (N=7), and Mexico City (N=1). For Vancouver participants, peripheral blood mononuclear cells (PBMCs) were isolated by density gradient separation and cryopreserved (−150°C, 90% Fetal Bovine Serum+ 10% DMSO). Participants recruited at other sites provided a leukapheresis sample from which PBMC were isolated and cryopreserved as above. All participants were on long-term, virally suppressive combination antiretroviral therapy (cART) at time of sampling, with the exception of one elite controller. Participants’ duration of untreated HIV infection ranged from 1 month to >10 years, though the exact duration was unknown for most participants. Ethical approval to conduct this study was obtained from the Institutional Review Boards of Simon Fraser University, Providence Health Care/University of British Columbia, Weill Cornell Medicine, and the George Washington University. All participants provided written informed consent.

### Quantitative Viral Outgrowth Assay (QVOA)

For participants for whom sufficient biological material was available, the Quantitative Viral Outgrowth Assay (QVOA) was performed as previously described^3^. Briefly, CD4+ T-cells were isolated from PBMCs by negative selection and plated in serial dilution at either 4 or 6 concentrations (12 wells/concentration, 24-well plates). CD4+ T-cells were stimulated with phytohemagglutinin (PHA, 2μg/mL) and irradiated allogeneic HIV-negative PBMCs were added to further induce viral reactivation. MOLT-4/CCR5 cells were added at 24hrs post-stimulation as targets for viral infection. Culture media (RPMI 1640 + 10% FBS + 1% Pen/Strep +50U/mL IL-2 + 10ng/mL IL-15) was changed every 3 days and p24 enzyme-linked immunosorbent assay (ELISA) was run on day 14 to identify virus-positive wells. Infectious Units per Million CD4+ T-cells (IUPM) was determined using the Extreme Limiting Dilution Analysis (ELDA) software (http://bioinf.wehi.edu.au/software/elda/)^13^. Culture supernatants from virus-positive wells were frozen (−80°C) for future use.

### Intact Proviral DNA Assay (IPDA)

Genomic DNA was isolated from a median 4.5 (interquartile range [IQR] 4-5) million CD4+ T-cells using the QIAamp DNA Mini Kit (Qiagen) with precautions to minimize DNA shearing. Intact HIV copies/million CD4+ T-cells were determined by droplet digital PCR (ddPCR) using the Intact Proviral DNA Assay (IPDA)^1^, where HIV and human RPP30 reactions were conducted independently in parallel and copies were normalized to the quantity of input DNA. In each ddPCR reaction, a median 7.5ng (IQR 7-7.5ng) (RPP30) or a median 750ng (IQR 700-750ng) (HIV) of genomic DNA was combined with ddPCR Supermix for Probes (no dUTPs, BioRad), primers (final concentration 900nM, Integrated DNA Technologies), probe(s) (final concentration 250nM, ThermoFisher Scientific) and nuclease free water. Primer and probe sequences (5’-> 3’) were: RPP30 Forward Primer-GATTTGGACCTGCGAGCG, RPP30 Probe-VIC-CTGACCTGAAGGCTCT-MGBNFQ, RPP30 Reverse Primer-GCGGCTGTCTCCACAAGT; RPP30 Shear Forward Primer-CCAATTTGCTGCTCCTTGGG, RPP30 Shear Probe-FAM-AAGGAGCAAGGTTCTATTGTAG-MGBNFQ, RPP30 Shear Reverse Primer-CATGCAAAGGAGGAAGCCG; HIV Ψ Forward Primer-CAGGACTCGGCTTGCTGAAG, HIV Ψ Probe-FAM-TTTTGGCGTACTCACCAGT-MGBNFQ, HIV Ψ Reverse Primer-GCACCCATCTCTCTCCTTCTAGC; HIV *env* Forward Primer-AGTGGTGCAGAGAGAAAAAAGAGC, HIV *env* Probe-VIC-CCTTGGGTTCTTGGGA-MGBNFQ, anti-Hypermutant *env* Probe-CCTTAGGTTCTTAGGAGC-MGBNFQ, HIV *env* Reverse Primer-GTCTGGCCTGTACCGTCAGC. For participant BC-004, an autologous *env* probe was designed by modifying the published IPDA *env* probe to match the participant’s sequence (VIC-CCTTGGGTTTCTGGGA-MGBNFQ). Droplets were prepared using either the Automated or QX200 Droplet Generator (BioRad) and cycled at 95°C for 10 minutes; 45 cycles of (94°C for 30 seconds, 59°C for 1 minute) and 98°C for 10 minutes, as previously described^1^. Droplets were analyzed on a QX200 Droplet Reader (BioRad) using QuantaSoft software (BioRad, version 1.7.4), where replicate wells were merged prior to analysis. Four technical replicates were performed for each participant sample, where a median (IQR) 243,100 (106,200-265,950) cells were assayed in total. Intact HIV copies (Ψ and *env* double-positive droplets) were corrected for DNA shearing based on the frequency of RPP30 and RPP30-Shear double positive droplets. The median (IQR) DNA shearing index (DSI), measuring the proportion of sheared DNA in a sample, was 0.31 (0.28-0.35), highly comparable to that reported by the original authors^1^. For the experiments that evaluated the IPDA’s ability to detect the specific *env* sequences harbored by participants 91C33 (Figure 2A) and OM5346 (Extended Data Figure 8B), synthetic templates (purified PCR amplicons for 91C33 and commercially-synthesized gBlocks [IDT] for OM5346) were used as targets. Total HIV *gag* copies/million CD4+ T-cells (Extended Data Figure 2) were measured as previously described^14^.

### Single-template, near-full length HIV genome proviral amplification and sequencing

Single-template, near-full-length proviral amplification was performed on DNA extracted from purified CD4+ T-cells by nested PCR using Platinum Taq DNA Polymerase High Fidelity (Invitrogen) as described^15^ such that ~25% of the resulting PCR reactions yielded an amplicon.

First round primers were: Forward - AAATCTCTAGCAGTGGCGCCCGAACAG, Reverse - TGAGGGATCTCTAGTTACCAGAGTC. Second round primers were: Forward - GCGCCCGAACAGGGACYTGAAARCGAAAG, Reverse-GCACTCAAGGCAAGCTTTATTGAGGCTTA. Reactions were cycled as follows: 92°C for 2 minutes; 10 cycles of (92°C for 10 seconds, 60°C for 30 seconds and 68°C for 10 minutes); 20 cycles of (92°C for 10 seconds, 55°C for 30 seconds and 68°C for 10 minutes); 68°C for 10 minutes^15,16^. Amplicons were sequenced using Illumina MiSeq technology and *de novo* assembled using an in-house modification of the Iterative Virus Assembler (IVA)^17^, using the custom software MiCall (https://github.com/cfe-lab/MiCall) or through collaboration with the Massachusetts General Hospital CCIB Core.

### *In silico* predicted IPDA results

Near full-length proviral sequences from participant BC-004 were used to predict IPDA results *in silico* under the assumption that proviruses with intact probe regions would be detected by the assay. The IPDA Ψ and *env* probe regions were excised from each proviral sequence and were considered defective if the probe region was absent or if it contained mutations associated with defective proviruses (*e.g*. those consistent with hypermutation^1^ or common splice donor site mutations^2^). All other sequences, regardless of whether they matched the published IPDA probe sequence, were considered intact in the probe region for the *in silico* reservoir composition assessment. A total of 381 proviruses were sequenced for BC-004, 338 of which contained at least one of the IPDA Ψ or *env* probe regions.

### Targeted sequencing of QVOA outgrowth viruses and participant-derived HIV sequences

For participant OM5346, HIV RNA was extracted from QVOA outgrowth viruses and pre-ART plasma, after which gp160 (QVOA) and gp120/Pol (plasma) were amplified from endpoint-diluted templates by nested RT-PCR using HIV-specific primers and high fidelity enzymes. For participants with suspected IPDA detection failure, and for whom HIV sequences were not already available, the IPDA Ψ and/or *env* amplicon regions were bulk-amplified from extracted proviral DNA using HIV-specific primers. Amplicons were sequenced using either Sanger (3730xl, Applied BioSystems) or Next Generation (Illumina MiSeq) technologies. Sanger chromatograms were analyzed using Sequencher (version 5.0.1, Gene Codes), while Illumina MiSeq reads were *de novo* assembled as described above.

### Antibody-dependent cellular cytotoxicity (ADCC) assays

Total CD4+ T-cells were isolated, as described above, from HIV-negative donors and activated with anti-CD3/anti-CD28 antibodies. Cells were then infected with the participant virus of interest collected from QVOA supernatant, and monitored by flow cytometry by intracellular staining for HIV-Gag (KC57-RD1, Beckman Coulter) until cells were >5% Gag+. CD4+ T-cells were washed and incubated with the broadly neutralizing antibody of interest (either 3BNC117 or 10-1074, 10μg/mL) for 2hrs. Natural Killer (NK) cells were negatively selected (EasySep, StemCell Technologies) from PBMCs of allogeneic, HIV-negative donors and activated using interleukin-15 (IL-15) provided by the National Cancer Institute Biological Research Branch. Activated NK cells were then co-cultured with infected, antibody-treated CD4+ T-cells for 16 hours at an effector-to-target (E:T) ratio of 1:1. Following co-culture, cells were stained with fluorophore-conjugated antibodies against human IgG, CD3, CD56, and CD4 (Biolegend), as well as intracellular Gag (KC57-RD1, Beckman Coulter) and LIVE/DEAD Fixable Aqua Stain amine-reactive dye (Invitrogen). The percentages (%) of infected cells remaining post-ADCC relative to no antibody control were then determined using the following formula: (1-((%Gag+ cells amongst viable CD3+ cells in no antibody condition)-(%Gag+ cells among viable CD3+ cells in test condition))/(%Gag+ cells amongst viable CD3+ cells under no antibody condition))*100.

### Phylogenetic Analysis of participant OM5346’s HIV RNA and reservoir diversity

For participant OM5346, single-genome HIV amplification and sequencing was performed from pre-ART plasma (sampled in 2012), proviruses from purified CD4+ T-cells isolated during suppressive cART (sampled in 2017 and 2019), and replication-competent HIV strains isolated by QVOA (from the 2017 sample) as described above. The 2012 and 2017/2019 samples were shipped directly to, and amplified in, two separate laboratories. A plasma HIV *pol* sequence derived from clinical drug resistance testing in 2012 was also incorporated into the analysis. *pol* and *gp120* sequences were specifically amplified or retrieved from near-full genome proviral sequences using GeneCutter (https://www.hiv.lanl.gov/content/sequence/GENE_CUTTER/cutter.html). Sequences were multiply-aligned using MAFFT (version 7.427)^18^ in a codon-aware manner and inspected in AliView^19^. Maximum-likelihood phylogenies were inferred using PhyML under a General Time Reversible nucleotide substitution model^20^ and visualized using FigTree (version 1.3.1).

### Design of Secondary *env* ddPCR primer/probe set

HIV-1 Subtype B *envelope* sequences, limited to one per participant, were retrieved from the HIV LANL database (N=4,670) and aligned. Sequence conservation was assessed across 16-25 base-pair intervals, and three intervals that maximized sequence conservation while meeting ddPCR assay specifications (*i.e*. amplicon length, T_m_) were chosen as secondary primer and probe locations. These were: Secondary *env* Forward Primer ACTATGGGCGCAGCGTC (representing nucleotides 7,809-7,825 in the HIV-1 genomic reference HXB2; predicted sequence conservation 79%), Secondary *env* Probe VIC-CTGGCCTGTACCGTCAG-MGBNFQ (HXB2 nucleotides 7,833-7,849, 83% conservation), Secondary *env* Reverse Primer CCCCAGACTGTGAGTTGCA (HXB2 nucleotides 7,939-7,921, 88% conservation). Reaction composition and cycling conditions were same as those used for the IPDA as described above. Accurate quantification using the Secondary *env* primer/probe set was verified using DNA extracted from the J-Lat 9.2 cell line (obtained from the NIH AIDS Reagent Program, Division of AIDS, NIAID, NIH, contributed by Dr. Eric Verdin)^21^ (Extended Data Figure 11).

### Statistical Analyses

All statistical analyses were performed using GraphPad Prism (version 8).

## Data and Sequence Availability

All sequences collected for the present study have been deposited in GenBank (Accession Numbers pending). Sequences for participant 91C33 were previously deposited under Accession Numbers: MH632930-MH632955^11^. All other data are available from the corresponding authors upon request, in compliance with institutional and REB requirements.

## Author Contributions Statement

NNK, YR, ZLB and RBJ designed the study. NNK, YR, WDCA, WD, PK, SHH, TMM, AS and AW performed experiments. NNK, YR, WDCA, WD, PK, SHH, TMM, AS, AW, DK, CJB, GQL and RML analyzed data. MH, CK, EB, MAO, PMDRE, AW, CC, WDH, LM, HG and ZLB provided participant samples. NNK, YR, ZLB and RBJ wrote the manuscript. All authors contributed to the critical revision of the manuscript.

## Acknowledgments

This work was supported by the Martin Delaney ‘BELIEVE’ Collaboratory (NIH grant 1UM1AI26617), which is supported by the following NIH Co-Funding and Participating Institutes and Centers: NIAID, NCI, NICHD, NHLBI, NIDA, NIMH, NIA, FIC, and OAR. It was also supported in part by the NIH funded R01 grants AI31798 and AI147845 (to BRJ) and a Canadian Institutes of Health Research (CIHR) project grant PJT-159625 (to ZLB). NNK is supported by a CIHR Vanier Doctoral Award. ZLB is supported by a Scholar Award from the Michael Smith Foundation for Health Research.

We thank the National Cancer Institute (NCI) Biological Research Branch for provision of IL-15 and IL-2, the NCI’s AIDS and Cancer Virus Program for provision of HIV p24 ELISA reagents, and the NIH AIDS Research and Reference Reagent Program for provision of the CCR5^+^ MOLT-4 cells. We thank Marina Caskey for providing de-identified samples from ART-naïve donors used in this study. We thank the BC Centre for Excellence in HIV/AIDS for support. We thank the Massachusetts General Hospital Center for Computational & Integrative Biology DNA Core, specifically Nicole Stange-Thomann, Amy Avery, Kristina Belanger, and Huajun Wang for providing us with the Illumina MiSeq deep sequencing service used in this manuscript. We thank Bruce Ganase, Jeff Knaggs, Daniel MacMillan and Hanwei Sudderuddin for research and bioinformatics support and Gursev Anmole, Mark Brockman and Robert Holt for helpful discussions. We thank Michel Nussenzweig, Christian Gaebler and Yotam Bar-On for provision of materials and helpful discussions. We gratefully acknowledge the contributions of the study participants, without whom this work would not be possible.

