## Extended Data Figures 1-11 for "HIV Diversity Considerations in the Application of the Intact Proviral DNA Assay (IPDA)"

Extended Data Figure 1

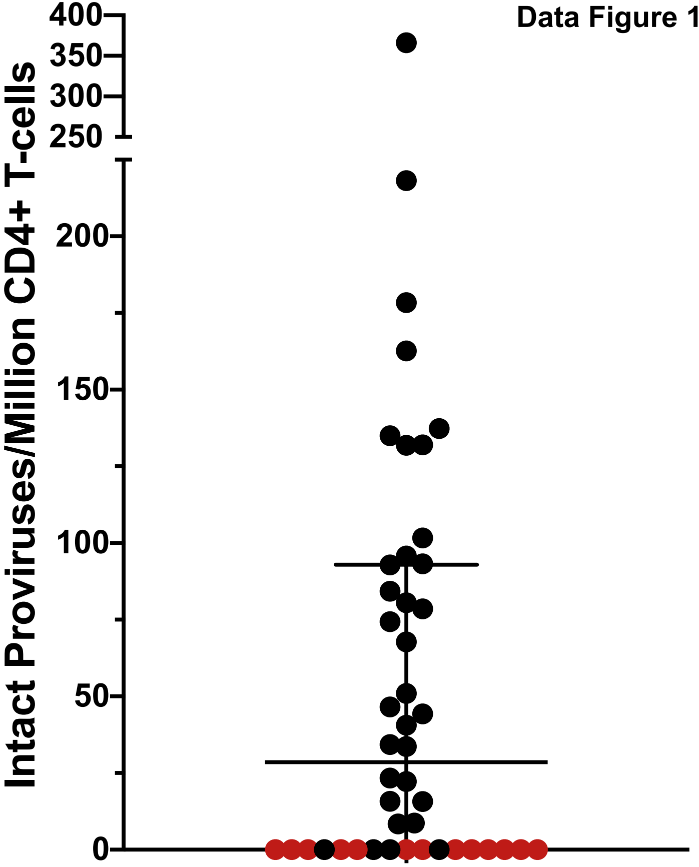

**Extended Data Figure 1: IPDA results for 46 study participants.** Line and error bars indicate cohort median and interquartile range; red datapoints represent 13 presumed instances of IPDA detection failure.

### Extended Data Figure 2

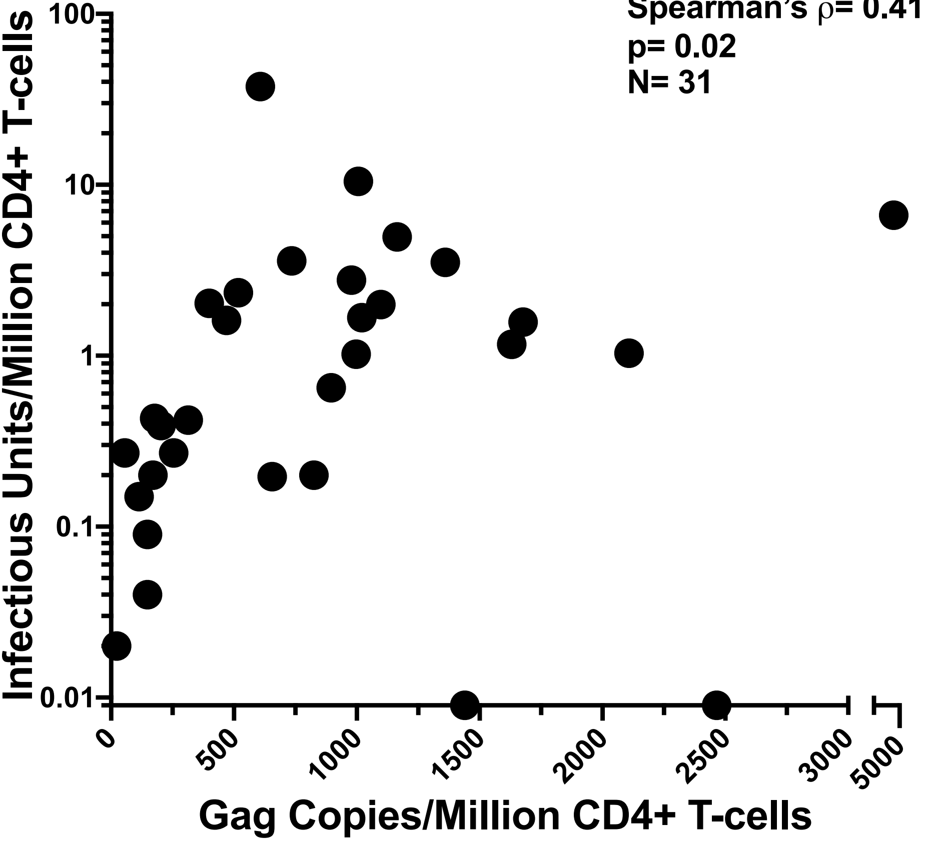

**Extended Data Figure 2: Spearman's correlation between QVOA and HIV *gag* copies/million CD4+ T-cells in a North American cohort.** Data were available for N=31 participants. Two individuals for whom no replication competent viruses were detected (IUPM= 0) are plotted on the X-axis.

**Extended Data Figure 3**

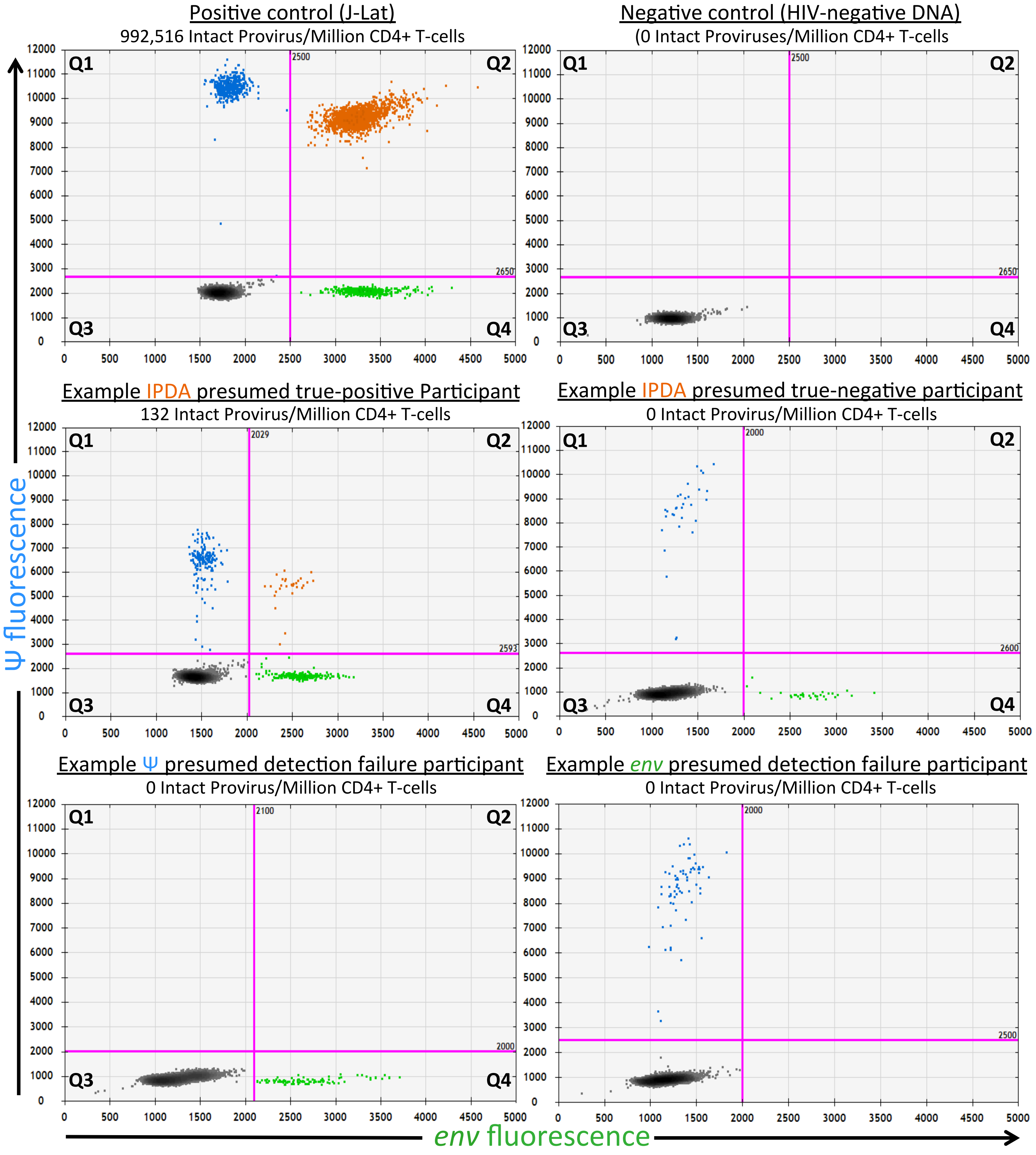

**Extended Data Figure 3: Example IPDA 2D plots for control and participant samples.** 2D ddPCR plots showing Ψ-single positive events (Q1, blue), Ψ- and *env*-double positive events (Q2, orange), double-negative events (Q3, grey) and *env*-single positive events (Q4, green), for positive control J-Lat cell line (top left; 1 copy of HIV per cell, droplets in Q1 and Q4 are a result of anticipated DNA shearing that occurs during DNA extraction that is subsequently corrected mathematically based on RPP30 shearing); HIV-negative donor negative control (top right); IPDA true-positive participant (middle left); IPDA true-negative participant (middle right) and IPDA detection failures (bottom row). Plots show merge of replicate wells.

Ext Data Figure 4

### Ψ sequence polymorphisms in IPDA Ψ detection failure samples

| ID | Ψ Forward Primer<br>CAGGACTCGGCTTGCTGAAG | Ψ Probe<br>ACTGGTGAGTACGCCAAAA | Ψ Reverse Primer<br>GCTAGAAGGAGAGAGATGGGTGC |
|---|---|---|---|
| HGLK001 | ------------------GC | ------------------T | ----------------------- |
| OM5365 | ------------------GC | R----------------TT | ----------------------- |
| WWH-012 | ------------------GC | ------------------- | ----------------------- |
| WWH-031 | ----------------A--- | -ACA--A----------TT | -------A------A--A----- |
| BC-014 | -------------------- | -AC---------------- | ----------------------- |

### *env* sequence polymorphisms in IPDA *env* detection failure samples

| ID | *env* Forward Primer<br>AGTGGTGCAGAGAGAAAAAAGAGC | *env* Probe<br>CCTTGGGTTCTTGGGA | *env* Reverse Primer<br>GCTGACGGTACAGGCCAGAC |
|---|---|---|---|
| HGLK002 | -----------------G------ | ---C--------ATC- | ------------------R- |
| HGLK005 | -----------*R-K---G------ | ---C------------ | ------------R------- |
| CIENI223 | ------------------------ | -----C---------- | ------------R------- |
| OM5334 | ---------------G-------- | ------A--------- | -------------R------ |
| CIRC0333 | ------------------------ | ---------T------ | -------------------A |
| WWH-011 | -----------------------G | ---------------- | -------------------- |
| WWH-026 | ------------------------ | ------T--------- | -------------------- |
| WWH-031 | ------------------------ | ---G-----TC----- | -------------------- |
| BC-004 | ---------A-------------- | ---------YC----- | -------------------- |

* = ATGCWR insertion prior to R

**Extended Data Figure 4: HIV polymorphism in IPDA primer/probe binding sites.** Primer and probe region sequences for participants with Ψ (top) or *env* (bottom) detection failure. Hyphens **(–)** indicate matches to the IPDA primer or probe; red letters indicate mismatches; asterisk indicates the location of an insertion.

Extended Data Figure 5

# 91C33

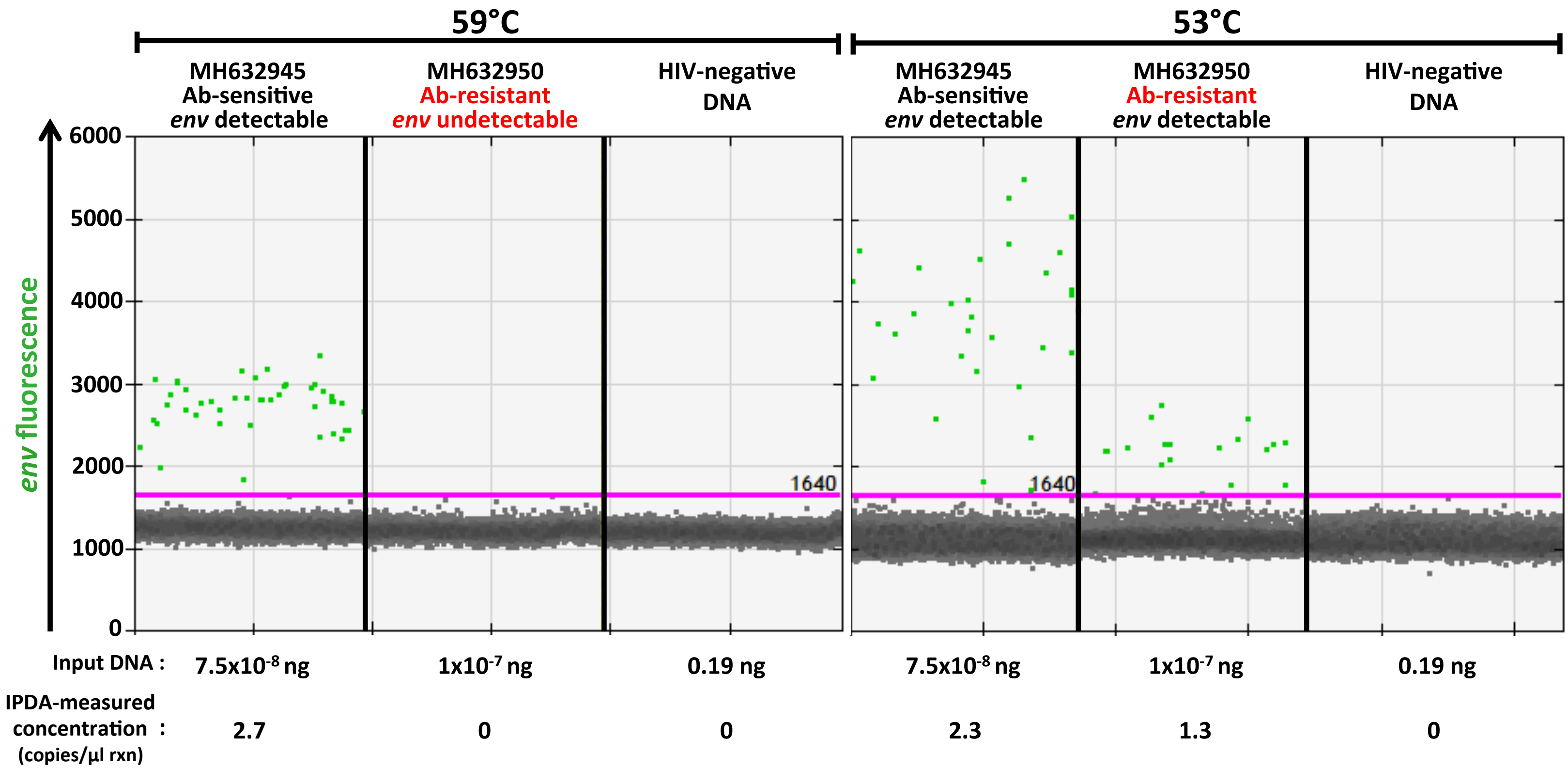

**Extended Data Figure 5: Modification of published IPDA conditions to reduce assay stringency allows for detection of 91C33 mismatch variant**. Representative ddPCR IPDA *env* plots for MH632945 (representing 91C33's bNAb-sensitive, IPDA *env* probe-matching HIV population) and MH632950 (representing 91C33's bNAb-resistant, IPDA env probe-mismatched HIV population); positive droplets are green and negative droplets are grey. Templates were purified *env* PCR products of equal length and comparable quantities. HIV-negative donor DNA served as a negative control. (left): results under published conditions; (right): result when assay annealing/extension temperature was reduced to 53°C.

Extended Data Figure 6A

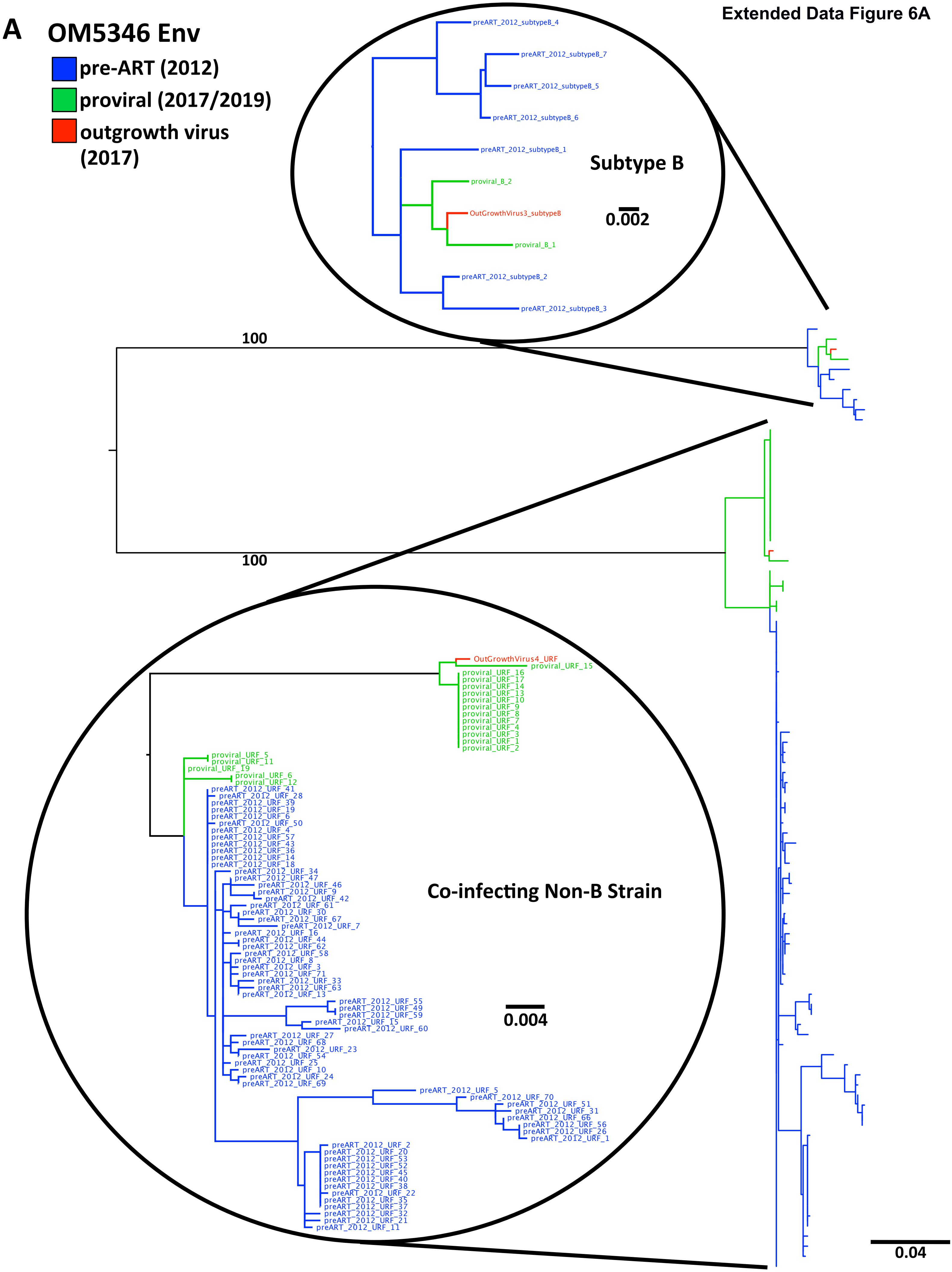

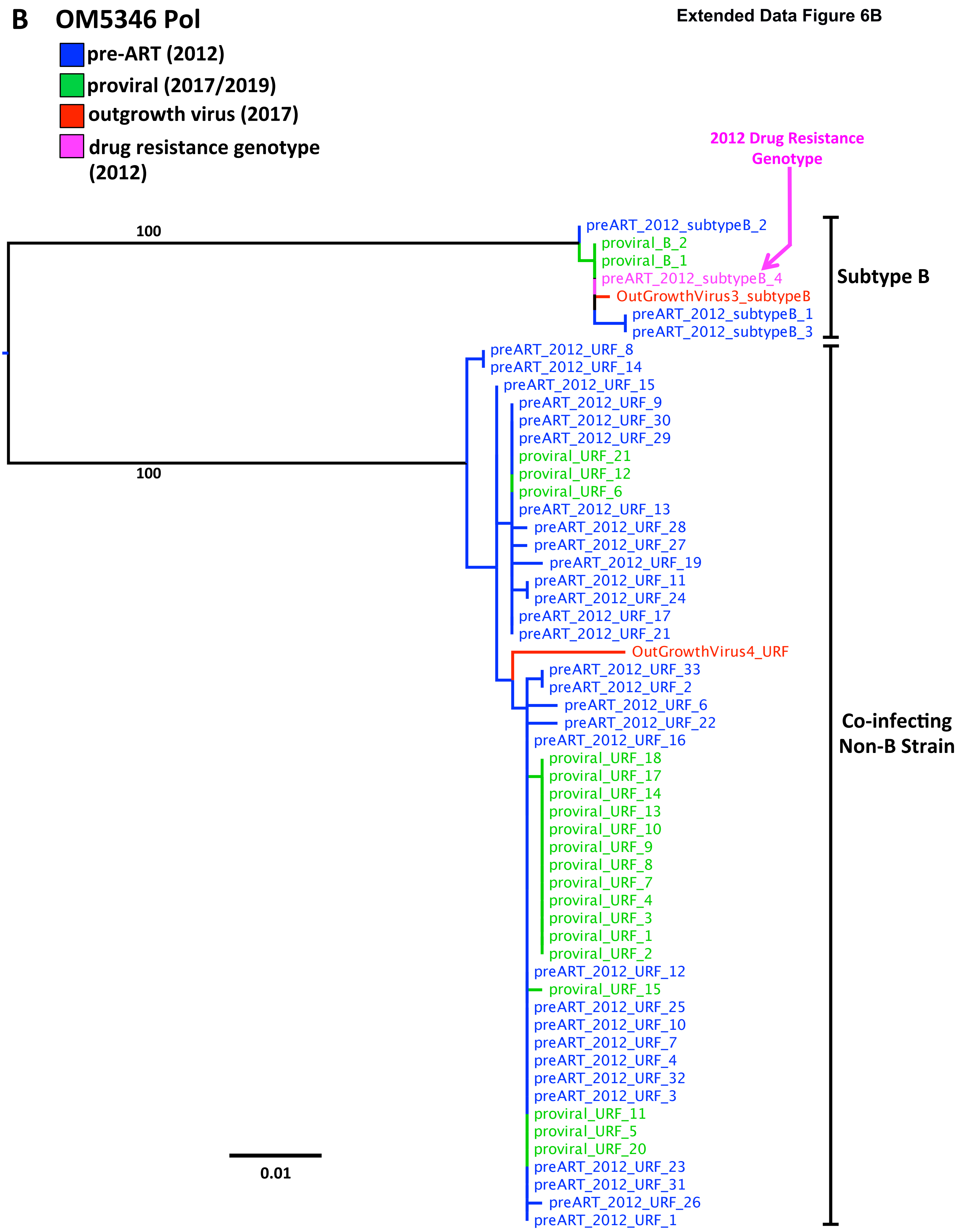
B
OM5346 Pol
pre-ART (2012)
proviral (2017/2019)
outgrowth virus (2017)
drug resistance genotype (2012)
2012 Drug Resistance Genotype
100
100
preART_2012_subtypeB_2
proviral_B_2
proviral_B_1
preART_2012_subtypeB_4
OutGrowthVirus3_subtypeB
preART_2012_subtypeB_1
preART_2012_subtypeB_3
Subtype B
preART_2012_URF_8
preART_2012_URF_14
preART_2012_URF_15
preART_2012_URF_9
preART_2012_URF_30
preART_2012_URF_29
proviral_URF_21
proviral_URF_12
proviral_URF_6
preART_2012_URF_13
preART_2012_URF_28
preART_2012_URF_27
preART_2012_URF_19
preART_2012_URF_11
preART_2012_URF_24
preART_2012_URF_17
preART_2012_URF_21
OutGrowthVirus4_URF
preART_2012_URF_33
preART_2012_URF_2
preART_2012_URF_6
preART_2012_URF_22
preART_2012_URF_16
proviral_URF_18
proviral_URF_17
proviral_URF_14
proviral_URF_13
proviral_URF_10
proviral_URF_9
proviral_URF_8
proviral_URF_7
proviral_URF_4
proviral_URF_3
proviral_URF_1
proviral_URF_2
preART_2012_URF_12
proviral_URF_15
preART_2012_URF_25
preART_2012_URF_10
preART_2012_URF_7
preART_2012_URF_4
preART_2012_URF_32
preART_2012_URF_3
proviral_URF_11
proviral_URF_5
proviral_URF_20
preART_2012_URF_23
preART_2012_URF_31
preART_2012_URF_26
preART_2012_URF_1
Co-infecting Non-B Strain
0.01

**Extended Data Figure 6: *Env* and *pol* phylogenies for participant OM5346, who is co-infected with two HIV strains. (A).** Maximum likelihood phylogeny inferred from single-genome-amplified *env* RNA sequences from pre-ART plasma (2012, blue), proviruses sampled on ART (2017 and 2019, green) and replication competent HIV sequences isolated from the reservoir (2017, red). **(B).** The *pol* phylogeny additionally includes a sequence from a clinical HIV drug resistance test performed in 2012 (pink). *Pol* sequences of replication-competent reservoir sequences (2017, red) were recovered from cells infected with these viruses *in vitro*. Scale bars indicate substitutions per nucleotide site. Numbers on main branches indicate branch support values.

### Extended Data Figure 7

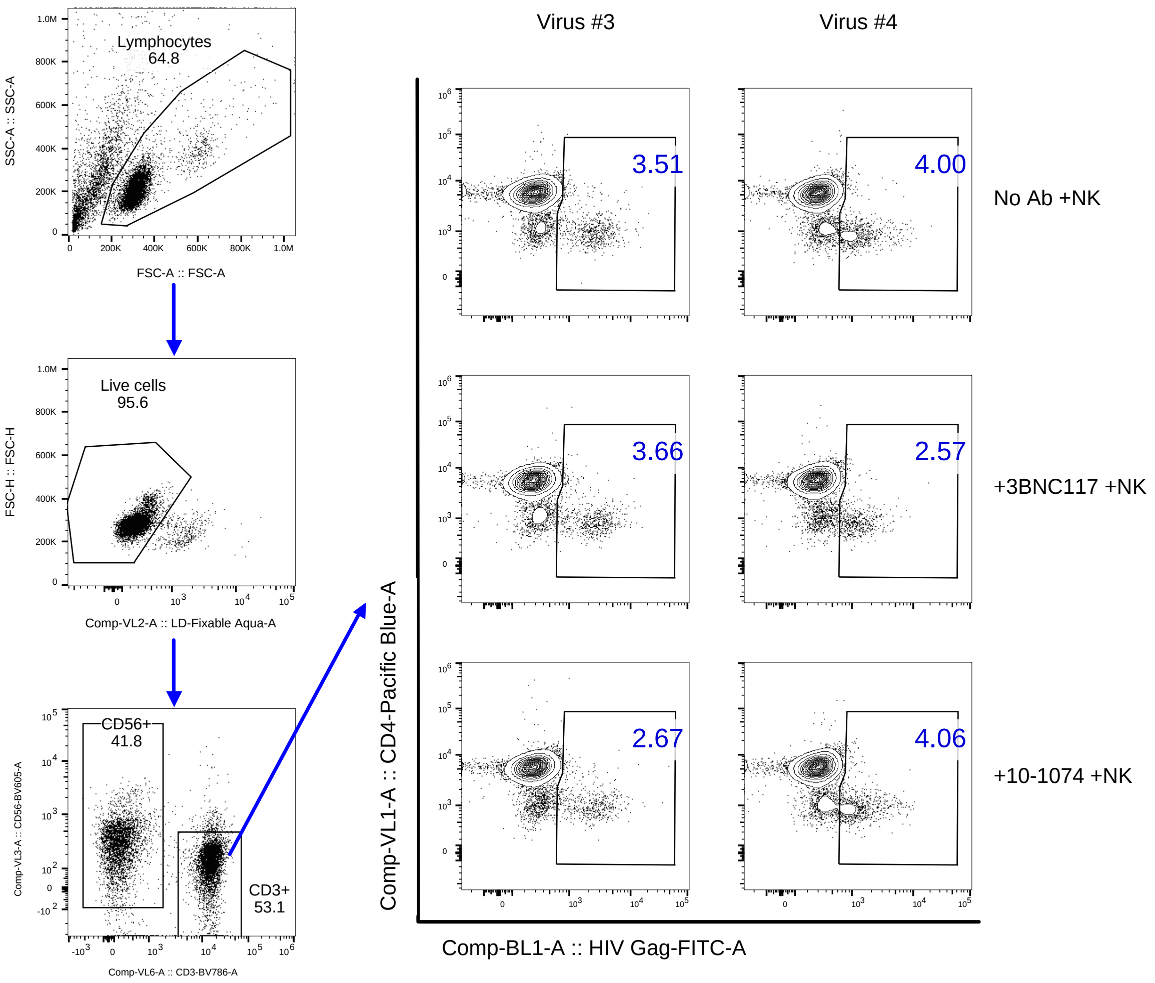

**Extended Data Figure 7: OM5346 ADCC gating strategy.** Representative flow plots showing flow cytometry gating strategy: Lymphocytes -> live cells (live/dead dye-) -> NK cells ($CD56^+$) and T-cells ($CD3^+$) -> HIV-positive cells (HIV $Gag^+$) from T-cell population. Activated $CD4^+$ T-cells from an HIV-negative donor were infected with OM5346 virus 3 (left) or virus 4 (right). NK cells were isolated from an HIV-negative donor. Plots on the top row show $CD4^+$ T-cells in control experiments (with no bNAb or NK cells added), 2nd row shows $CD4^+$ T cells remaining after addition of NK cells only, 3rd row shows $CD4^+$ T cells remaining after addition of 3BNC117 and NK cells, 4th row shows $CD4^+$ T cells remaining after addition of 10-1074 and NK cells. CD4 downregulation in HIV-positive cells (HIV $Gag^+$) occurs due to Nef-induced CD4 downregulation.

**A** Ext Data Figure 8

### Virus-infected cells: No detection of OM5346 Virus 4 by IPDA

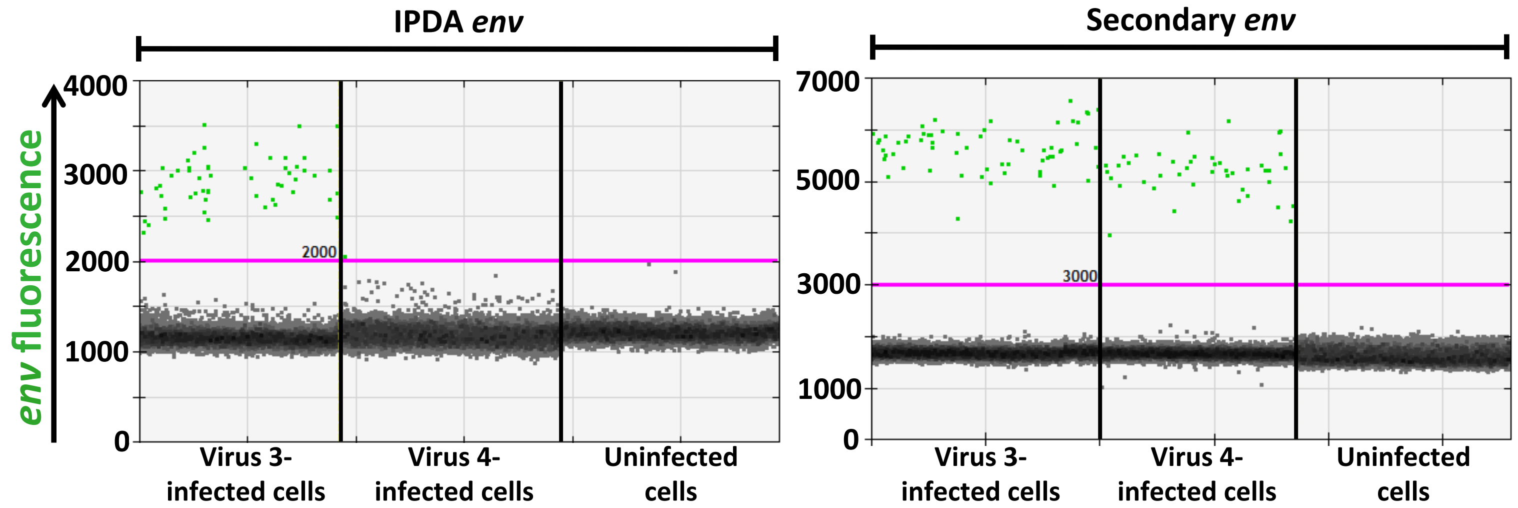


| | | | | | | | |
|---|---|---|---|---|---|---|---|
| input DNA | 0.75 ng | 0.75 ng | 0.75 ng | | 0.75 ng | 0.75 ng | 0.75 ng |
| IPDA-measured concentration (copies/µl rxn) | 3.6 | 0.06 | 0 | | 3.6 | 3.1 | 0 |

**B**

### Detection of Virus 4 pure template at high, but not participant-*like*, concentrations

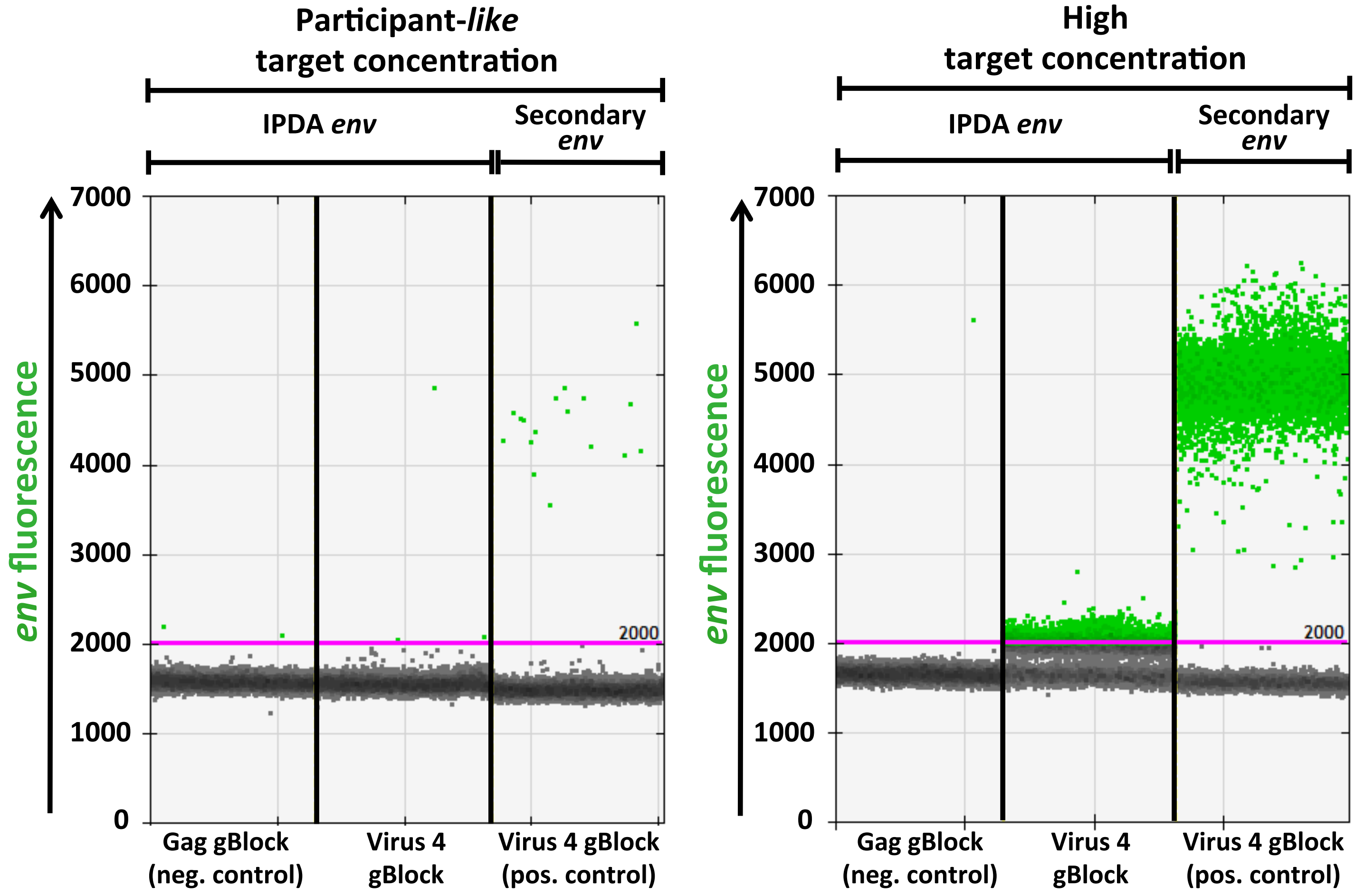


| | | | | | | | |
|---|---|---|---|---|---|---|---|
| input DNA (nmol) | $1x10^{-6}$ | $1x10^{-6}$ | $1x10^{-6}$ | | $6.7x10^{-4}$ | $6.7x10^{-4}$ | $6.7x10^{-4}$ |
| IPDA-measured concentration (copies/µl rxn) | 0.18 | 0.25 | 1.5 | | 0.09 | 456 | 914 |

**Extended Data Figure 8: OM5346 Virus 4 sequence is undetectable by the IPDA at participant-like concentrations. (A).** Representative 1D *env* plots using IPDA (left) or the secondary *env* reaction (right), from CD4$^+$ T-cells infected with OM5346 virus 3 (IPDA *env* probe match), virus 4 (IPDA *env* probe G13A mismatch) or uninfected cells (negative control). Positive droplets are green; negative droplets are grey. Copies/µl reaction as calculated from the experimental data are shown below the plots. The left panel is the same as Figure 2B. **(B).** Representative 1D *env* plots of OM5346 virus 4, tested as a synthetic DNA gene fragment ("Virus 4 gBlock"), using IPDA and secondary *env* reactions, at different input concentrations. A Gag synthetic gene fragment ("Gag gBlock") served as the negative control. Input DNA quantity and copies/µl reaction as calculated from the experimental data are shown below the plot. These observations provide a possible explanation to reconcile the original authors' ability to detect this sequence using a pure plasmid template and our inability to detect it at conditions mimicking a participant sample.

Extended Data Figure 9

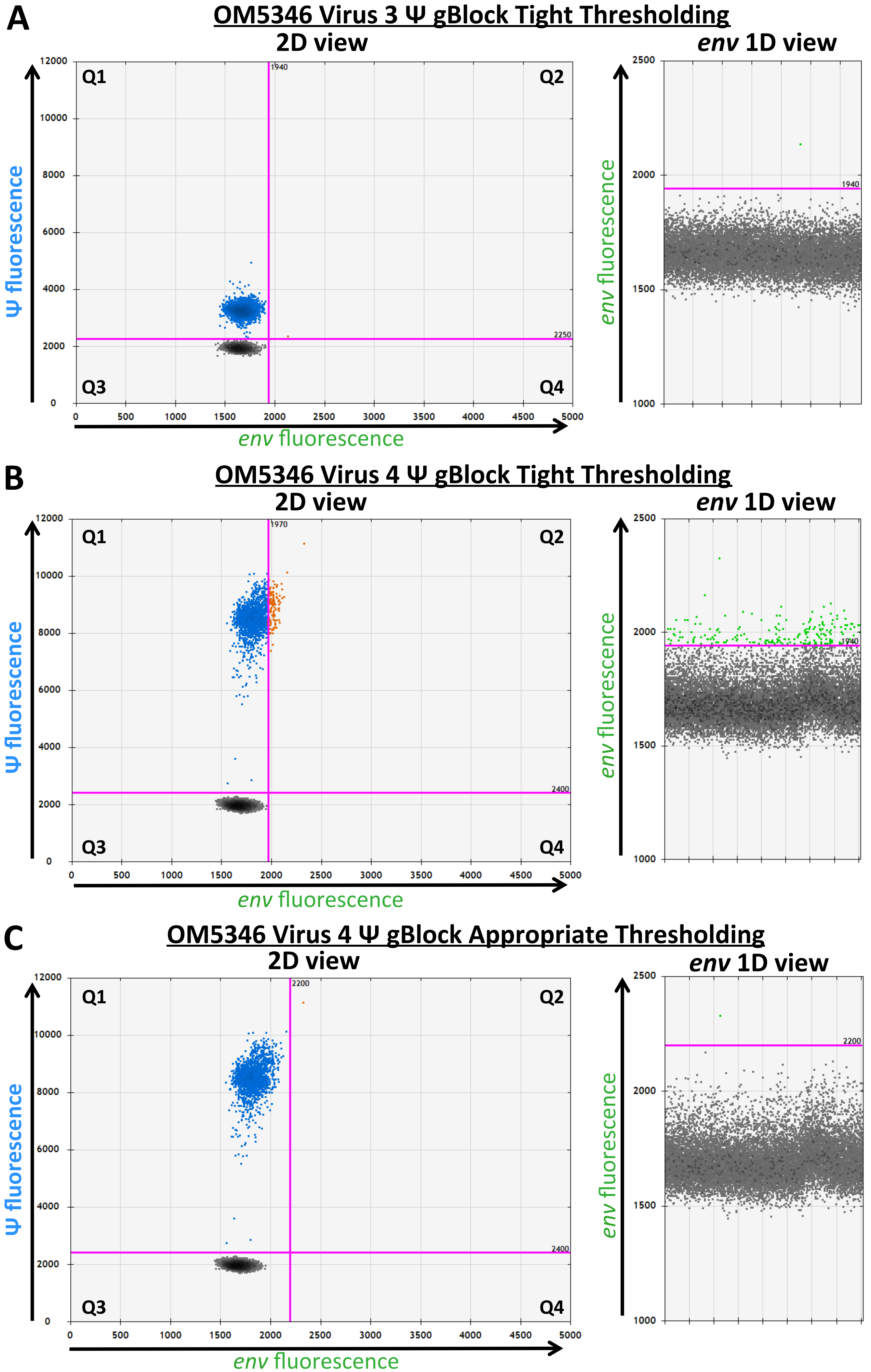

**Extended Data Figure 9: Sequence-specific fluorescence spillover prohibits tight thresholding on negative populations in the IPDA.** **(A)** 2D (left) and *env* 1D (right) plots of OM5346 virus 3 Ψ region sequence, tested as a synthetic DNA gene fragment ("Virus 3 Ψ gBlock") without corresponding *env* template. Minimal Ψ- to *env*- channel spillover occurs when drawing the positive droplet threshold tightly to the double negative population (note the presence of one false-double positive droplet). **(B)** 2D (left) and *env* 1D (right) plots of OM5346 virus 4 Ψ region sequence, tested as a synthetic DNA gene fragment ("Virus 4 Ψ gBlock"), without corresponding *env* template. With this sequence, drawing of a tight threshold yields marked spillover of Ψ (FAM) fluorescence into the *env* (VIC) channel, yielding false-positive *env* (and by extension, false-positive intact) signal. **(C)** 2D (left) and *env* 1D (right) plots of OM5346 Virus 4 Ψ region sequence, tested as a synthetic DNA gene fragment ("Virus 4 Ψ gBlock") without corresponding *env* template, with a threshold drawn at an appropriate distance from the double negative population. This threshold accommodates the sequence-specific shift in the Ψ-positive to avoid the creation of false-positive intact or *env*-positive droplet population (note the presence of a single false-double positive droplet).

Ext Data Figure 10

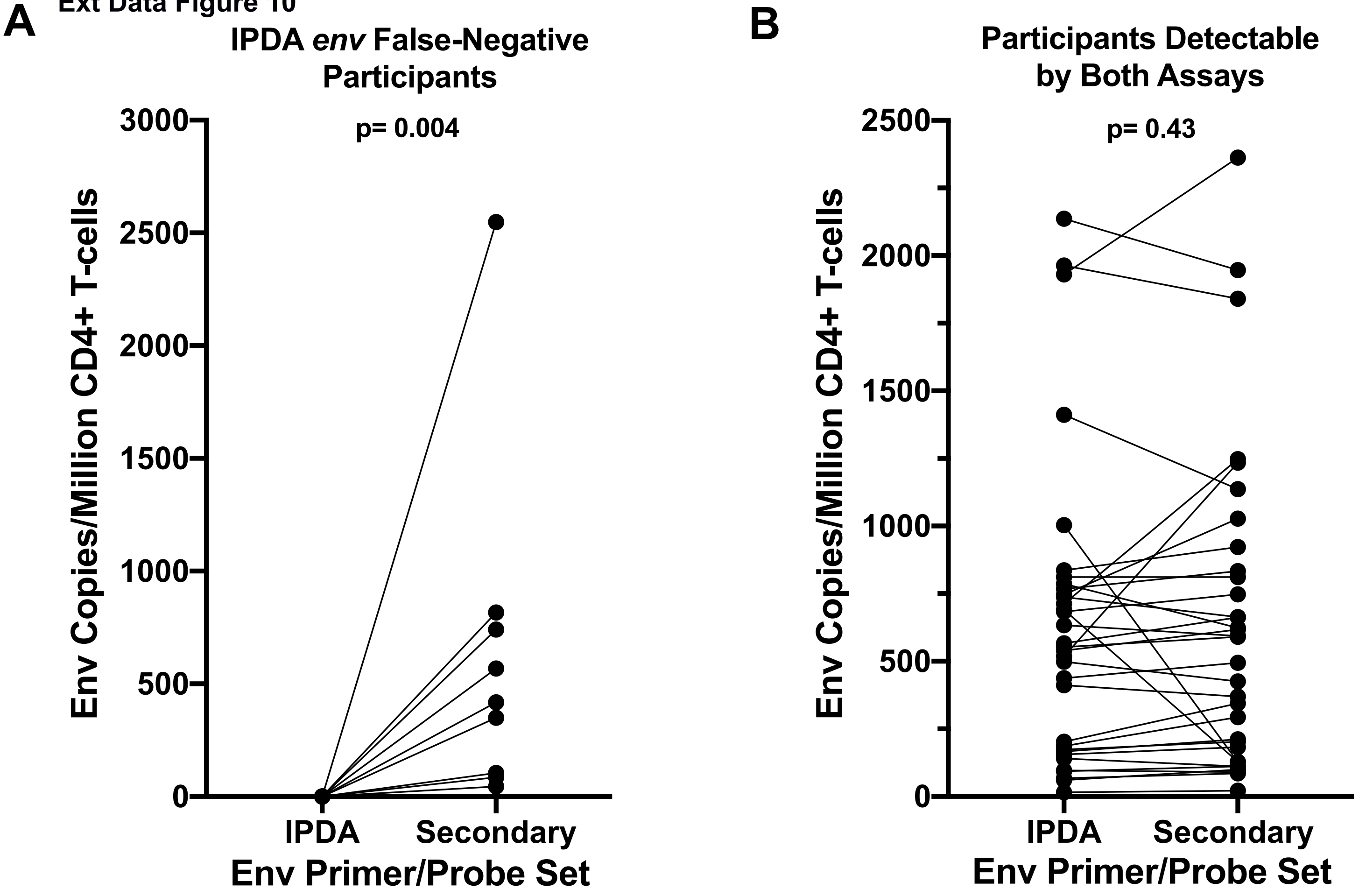

**Extended Data Figure 10: Performance of the secondary *env* primer/probe set. (A)** A secondary *env* primer/probe set rescued detection in all 9 cases of IPDA *env* detection failure **(B)** In 33 participants whose reservoir was detectable by both IPDA and secondary *env* primer/probe sets, measurements showed no significant differences by Wilcoxon signed-rank test.

### Ext Data Figure 11

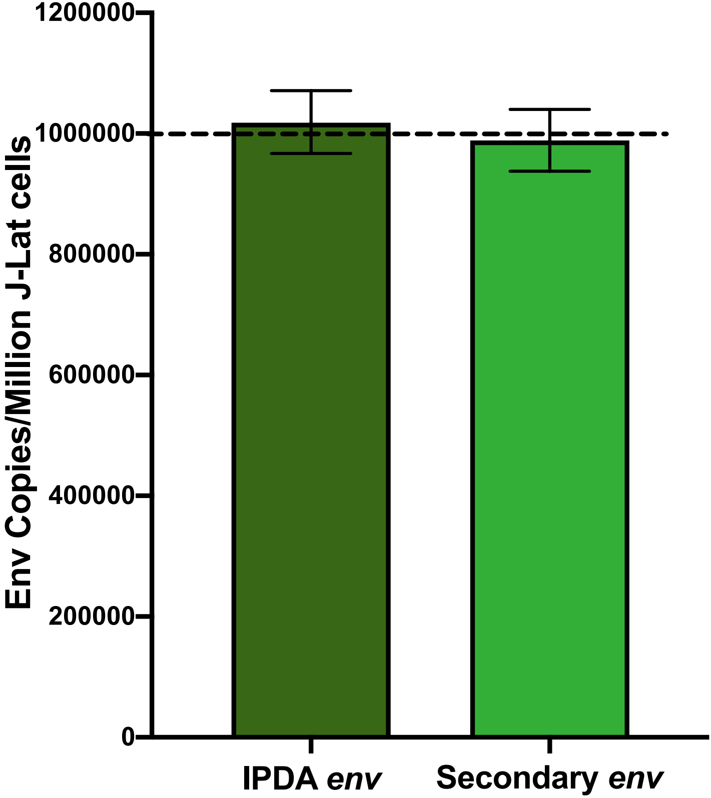

**Extended Data Figure 11: Comparable detection of expected 1:1 HIV-to-Cell ratio in J-Lat cells by IPDA and Secondary *env* reactions.** Error bars indicate 95% total Poisson confidence interval.
